# Resistance to antibacterial peptide nucleic acids through altered ribosome function

**DOI:** 10.64898/2026.09.05.749517

**Authors:** Jakob Frimodt-Møller, Irene M. Román, Thomas Prossliner, Lotta J. Happonen, Karen L. Nielsen, Susanne Häußler, Elio Rossi, Michael O’Connor, Peter E. Nielsen, Anders Løbner-Olesen

**Affiliations:** Department of Biology, University of Copenhagen, Denmark; Department of Clinical Sciences, Division of Infection Medicine, Lund University, SE-22184 Lund, Sweden; Structural Proteomics, Science for Life Laboratory, Lund University, SE-22184 Lund, Sweden; Department of Clinical Microbiology, Rigshospitalet, Denmark; Department of Biosciences, University of Milan, Italy; School of Science and Engineering, University of Missouri-Kansas City, USA; Department of Cellular and Molecular Medicine, University of Copenhagen, Denmark

## Abstract

When used as antibacterial agents, Peptide Nucleic Acid (PNAs) are generally designed to base-pair with complementary sequences of an essential mRNA and block translation initiation. Although bacterial susceptibility to peptide-conjugated PNAs is strongly influenced by cellular uptake, intracellular determinants of PNA activity remain poorly understood. Here, we identify a ribosome-centered mechanism of resistance to antibacterial PNAs in *Escherichia coli*. The *rpsL*_*I82N*_ mutation increased the minimum inhibitory concentration (MIC) of an arginine-rich cell penetrating peptide-conjugated PNA targeting *acpP* eightfold. The effect of *rpsL*_*I82N*_ was additive with mutations that reduce PNA entry into bacterial cells

PNA resistance conferred by *rpsL*_*I82N*_ was independent of carrier peptide and also applied when naked PNA was tested in an envelope-permeable strain. Similarly, *rpsL*_*I82N*_-associated resistance was independent of the targeted mRNA, as it applied to PNAs targeting either the Shine–Dalgarno or AUG region of *acpP* or *ftsZ* mRNA. Several additional substitutions within residues 74–82 of ribosomal protein uS12 conferred PNA resistance. Because resistance occurred among both error-restrictive and ribosomal-ambiguity alleles, it did not correlate with the classical decoding-fidelity phenotypes measured by stop-codon readthrough.

Proteomic analysis revealed widespread changes in proteins under post-transcriptional control in *rpsL*_*I82N*_ cells. The effect on selected sRNA-regulated genes correlated with the location of the sRNA-binding site: repression was less efficient when binding overlapped the translation-initiation region whereas it was more efficient when regulatory sites were located outside this region. In parallel, *rpsL*_*I82N*_ increased the 30S and 50S ribosomal fractions and reduced the 70S fraction. Both loss of KsgA, which disrupts 30S maturation, and treatment with kasugamycin, which perturbs translation initiation, increased PNA resistance.

We propose that *rpsL*_*I82N*_ alters uS12-dependent 30S assembly and initiation-complex dynamics, thereby changing the effective occupancy of mRNA translation-initiation regions. This limits access of PNAs and sRNAs to overlapping sequences, resulting in PNA resistance and reduced sRNA-mediated repression. Conversely, a longer-lived 30S initiation state may facilitate sRNA binding to flanking regions and strengthen repression. These findings identify the ribosome–mRNA interface as an intracellular determinant of antibacterial PNA susceptibility.

## Introduction

Peptide nucleic acids are synthetic nucleic-acid analogues in which the phosphodiester backbone of DNA or RNA is replaced by an uncharged pseudo-peptide backbone. PNAs bind complementary RNA with high affinity and sequence specificity and are resistant to degradation by cellular nucleases and proteases. In bacteria, antibacterial PNAs are generally designed to target the translation-initiation region of an essential mRNA. Hybridization across the Shine–Dalgarno sequence, start codon or intervening region sterically prevents productive ribosome engagement and inhibits synthesis of the encoded protein. Early studies established that peptide-conjugated PNAs targeting the essential *acpP* transcript could inhibit *E. coli* growth and that sequences surrounding the translation start codon were particularly susceptible to antisense inhibition [1,2]. More recent cell-free analyses have confirmed that PNAs directly inhibit translation of their intended targets and have helped define the sequence-complementarity requirements governing on- and off-target translation inhibition [3–5].

Efficient delivery remains a major limitation of antibacterial PNA activity. Because naked PNA penetrates the Gram-negative envelope poorly, PNAs are commonly conjugated to cationic or amphiphilic bacterial-penetrating peptides. Different carriers encounter different envelope barriers and use distinct routes to reach the cytoplasm. Uptake of shorter KFF-derived conjugates can depend on the inner-membrane protein SbmA, whereas arginine-rich RXR conjugates are strongly influenced by membrane potential and envelope-stress physiology [6–8]. The outer lipopolysaccharide layer also affects susceptibility in a carrier-dependent manner: disruption of the LPS outer core preferentially increases susceptibility to KFF–PNA while having much less effect on arginine-rich conjugates [9]. These findings established impaired delivery as a major route to PNA resistance but also suggested that “PNA susceptibility” represents the combined outcome of carrier chemistry, envelope genotype and intra-cellular antisense activity.

We previously selected *E. coli* mutants with increased resistance to an RXR-conjugated PNA targeting the AUG region of the essential *acpP* transcript. The most resistant isolate carried the *cpxR*_*L20Q*_ mutation, insertion-sequence disruptions of *waaB* and *waaO*, and a *rpsL*_*I82N*_ mutation in the gene encoding ribosomal protein uS12. The *cpxR*_*L20Q*_ mutation led to activation of the Cpx envelope-stress response, which reduced the activity of the arginine-rich PNA conjugate by lowering respiration and membrane potential, but only accounted for part of the resistance. This left open the possibility that PNA activity could also be limited after the conjugate had entered the cytoplasm [8].

Transcript abundance does not reliably predict whether an essential gene will be a vulnerable antibacterial PNA target, and predicted PNA–RNA melting temperature correlates better with inhibition in purified translation systems than with whole-cell antibacterial activity. PNA-induced depletion of the target mRNA is also variable and is not required for growth inhibition. Global RNA profiling has further shown that peptide identity and target sequence produce distinct physiological responses in addition to the intended target knockdown [3,4]. These observations indicate that intracellular PNA efficacy depends on more than hybridization thermodynamics and target abundance. The structural and kinetic state of the target mRNA, including its occupancy by ribosomes and RNA-binding proteins, is likely to contribute to the fraction of transcripts that can be captured by PNA.

Translation-initiation regions are also major targets of endogenous small regulatory RNAs. Base-pairing sRNAs, often assisted by Hfq, regulate translation and mRNA stability by interacting with short regions of target transcripts. Many negatively acting sRNAs bind across the Shine–Dalgarno sequence or start codons and directly compete with the 30S subunit. A well-characterized example is MicA-mediated repression of *ompA* in which toeprinting demonstrated interference with ribosome binding at the translation-initiation region [10]. Other sRNAs bind upstream of the canonical ribosome-binding site or within the early coding sequence and can inhibit translation through RNA restructuring, Hfq-dependent mechanisms, interference with initiation-complex formation or recruitment of RNA-degradation machinery [11]. The recent expansion of the experimentally supported GcvB regulon to more than 50 direct targets further illustrates the diversity of binding-site positions and regulatory mechanisms used by a single bacterial sRNA [12].

Translation initiation is a multistep process in which the 30S ribosomal subunit, mRNA, initiator tRNA and initiation factors form a pre-initiation complex that subsequently matures into a 30S initiation complex that covers the Shine-Dalgarno (SD) sequence and the start codon. Release or rearrangement of initiation factors permits 50S joining and formation of an elongation-competent 70S ribosome. Changes in the abundance, structure or lifetime of these intermediates could alter how long the translation-initiation region remains accessible to a PNA or sRNA. This is demonstrated by recent structural work showing that kasugamycin and GE81112 arrest distinct stages of 30S initiation-complex formation, emphasizing that translation initiation comprises of kinetically and structurally separable states rather than a single ribosome-binding event [13].

Ribosomal protein uS12, encoded by *rpsL*, is positioned within the 30S decoding centre and participates in the conformational network governing tRNA selection. Classical *rpsL* mutations can confer streptomycin resistance and produce either error-restrictive, hyper-accurate ribosomes or error-prone ribosomal-ambiguity phenotypes. However, mutational analyses have shown that substitutions throughout S12 can alter fidelity without necessarily changing streptomycin susceptibility [14]. Beyond these classical roles, uS12 has been implicated in RNA folding, ribosome assembly and translation initiation. Purified *E. coli* S12 can act as a broad-specificity RNA chaperone in vitro [15], and single-molecule experiments showed that S12 promotes cotranscriptional pre-16S rRNA folding and stable S4 recruitment during 30S assembly [16]. Genetic studies have also linked uS12 substitutions to IF3-dependent initiator-tRNA selection [17], while other alterations in S12 have been associated with defects in small-subunit assembly, initiation, elongation and recycling [18,19]. Thus, an S12 substitution could influence PNA activity by altering rRNA folding, ribosome assembly or initiation-state dynamics independently of its effect on classical decoding accuracy.

Late 30S maturation provides an additional connection between small-subunit structure and initiation. KsgA dimethylates A1518 and A1519 in helix 45 of 16S rRNA and functions as a late-stage ribosome-assembly factor. Structural studies show that KsgA binding is incompatible with mature positioning of helix 44 and thereby couples completion of small-subunit assembly to formation of the decoding site and productive subunit joining [20,21]. Perturbing KsgA-dependent maturation can consequently alter the distribution and functional state of 30S particles, potentially changing the competition between ribosomes and antisense molecules at the mRNA translation-initiation region.

Here, we identify *rpsL*_*I82N*_ as the second major PNA-resistance determinant in the evolved *E. coli* strain Evo-3. We define the breadth of this phenotype across peptide carriers, target transcripts and translation-initiation sites, and examine its relationship to S12 variation, fitness, translation fidelity, sRNA regulation and ribosome-state distribution. Our findings identify an intracellular resistance mechanism in which altered 30S occupancy of the translation-initiation region affects access by both PNAs and endogenous sRNAs.

## Results

### An altered ribosome confers resistance to an antisense PNA

We previously identified a strain (Evo-3), carrying the mutations *cpxR*_*L20Q*_, *rpsL*_*I82N*_, IS1:*waaB*, IS1:*waaO*, as resistant to an RXR-conjugated PNA targeting the AUG start-codon region of the essential *acpP* mRNA [8]. The *cpxR*_*L20Q*_ mutation is responsible for reduced RXR-PNA uptake but only partly responsible for the resistance (Table 1) [8]. We therefore proceeded to determine which of the other mutations present in Evo-3 contributed to RXR-PNA resistance.

**Table 1.** Values indicate MICs unless otherwise stated. RXR–PNA targets the *acpP* AUG/start-codon region unless indicated. RXR–PNA_SD_ targets the *acpP* Shine– Dalgarno region. RXR–PNA_*ftsZ*_ targets the *ftsZ* translation-initiation region. Naked PNA was tested in the AS19 background. “—” indicates not determined.^a^indicates a genotype otherwise as wild-type. ^b^ indicates nonsynonymous *rpsL* mutants isolated at the Dept. of Clinical Microbiology, Rigshospitalet, Denmark. Doubling time was determined in MHB II at 37°C.

| Strain | Rel. genotype | RXR-PNA<br>(μM) | RXR-PNA<sub>SD</sub><br>(μM) | RXR-PNA<sub>ftsZ</sub><br>(μM) | KFF-PNA<br>(μM) | PNA<br>(μM) | Strep<br>(μg/mL) | Doubling<br>time (min) |
| --- | --- | --- | --- | --- | --- | --- | --- | --- |
| Wild-type |  | 0.5 | 4 | 0.5 | 0.5 | >16 | 4 | 24 |
| Evo-3 | *cpXR<sub>L20Q</sub>*, *rpsL<sub>I82N</sub>*,<br>*IS1:waaB*,<br>*IS1:waaO* | 8 | - | - | 0.5 | - | 4 | 37 |
| *cpXR<sub>L20Q</sub>*<sup>a</sup> |  | 4 | - | - | - | - | - | - |
| *rpsL<sub>I82N</sub>*<sup>a</sup> |  | 4 | >16 | 4 | 2 | - | 4 | 33 |
| *rpsL<sub>I82N</sub> cpXR<sub>L20Q</sub>*<sup>a</sup> |  | 8 | - | - | - | - | - | - |
| ΔwaaBO<sup>a</sup> | | 0.5 | - | - | 0.25 | - | - | - |
| *rpsL<sub>I82N</sub> ΔwaaBO<sup>a</sup>* |  | 4 | - | - | 0.5 | - | - | - |
| Evo-3 pCA24n |  | 8 | - | - | - | - | - | 37 |
| Evo-3 pCA24n:*rpsL* |  | 4 | - | - | - | - | - | 23 |
| AS19 |  | - | - | - | - | 0.25 | - | - |
| AS19 pALO277 |  | - | - | - | - | 0.25 | - | - |
| AS19 pALO277:*rpsL* |  | - | - | - | - | 0.25 | - | - |
| AS19 pALO277:*rpsL<sub>I82N</sub>* |  | - | - | - | - | 2 | - | - |
| *rpsL<sub>K44I</sub>*<sup>a</sup> |  | 0.5 | - | - | - | - | - | - |
| *rpsL<sub>L74P</sub>*<sup>a</sup> |  | 2 | - | - | - | - | - | - |
| *rpsL<sub>S78C</sub>*<sup>a</sup> |  | 0.5 | - | - | - | - | - | - |
| *rpsL<sub>V79E</sub>*<sup>a</sup> |  | 0.5 | - | - | - | - | - | - |
| *rpsL<sub>I80N</sub>*<sup>a</sup> |  | 2 | - | - | - | - | - | - |
| *rpsL<sub>L81R</sub>*<sup>a</sup> |  | 2 | - | - | - | - | - | - |
| *rpsL<sub>I82F</sub>*<sup>a</sup> |  | 4 | - | - | - | - | - | 41 |
| *rpsL<sub>R86H</sub>*<sup>a</sup> |  | 0.5 | - | - | - | - | - | - |
| *rpsL<sub>R86S</sub>*<sup>a</sup> |  | 0.5 | - | - | - | - | - | - |
| *rpsL<sub>T64A</sub>*<sup>b</sup> |  | 0.25 | - | - | - | - | - | - |
| *rpsL<sub>I80L</sub>*<sup>b</sup> |  | 0.5 | - | - | - | - | - | - |
| *rpsL<sub>I82N</sub>*<sup>b</sup> |  | 4 | - | - | - | - | - | - |

Loss of WaaB and WaaO, resulting in loss of the outer core of the LPS layer of the outer membrane, had no effect on resistance to RXR-PNA (Table 1). This agrees with our previous finding that the inner membrane is the main barrier for entry of the RXR-PNA conjugate [8]. We subsequently examined the *rpsL*_*I82N*_ mutation, which results in replacement of isoleucine with asparagine at position 82 in the S12 protein of the 30S ribosomal subunit. When introduced into wild-type cells, *rpsL*_*I82N*_ conferred an eight-fold increase in resistance to RXR-PNA compared with the wild type, with the MIC increasing from 0.5 to 4 μM. This was comparable to the resistance conferred by *cpxR*_*L20Q*_ alone (Table 1).

When *rpsL*^I82N^ was combined with *cpxR*^L20Q^ in otherwise wild-type cells, the MIC of RXR-PNA increased from 4 to 8 μM (Table 1). Thus, the resistance observed in Evo-3 can be explained by two effects: reduced translocation resulting from the CpxR_L20Q_ or reduced intracellular activity mediated by RpsL_I82N_. In agreement with this interpretation, expression of wild-type RpsL in Evo-3 partially reversed resistance from 8 to 4 μM, corresponding to the level conferred by *cpxR*_L20Q_ alone (Table 1). These results identify *cpxR*_L20Q_ and *rpsL*_*I82N*_ mutations as the main drivers of RXR-PNA resistance.

To test whether *rpsL*_*I82N*_ conferred resistance to the PNA component rather than specifically to the RXR peptide carrier, the same anti-*acpP* PNA was conjugated to a KFF carrier. The *rpsL*_*I82N*_ mutant showed a two-fold increase in resistance to KFF-PNA compared with the wild type (Table 1). We next tested naked PNA in the envelope-permeable *E. coli* strain AS19 [22]. Wild-type and mutant *rpsL* alleles were cloned into the low-copy F-based plasmid pALO277, generating pALO277:*rpsL* and pALO277: *rpsL*_*I82N*_, respectively. The native pALO277 plasmid has approximately one to two copies per genome equivalent [23,24], which is equal to or slightly higher than the copy number of *rpsL* at its chromosomal location. AS19, AS19-pALO277, and AS19-pALO277:*rpsL* remained sensitive to naked PNA, whereas AS19-pALO277: *rpsL*_*I82N*_ showed a four-fold increase in resistance (Table 1). These findings demonstrate that *rpsL*_*I82N*_ mediates resistance independently of the peptide carrier and suggest that it acts through an intracellular mechanism.

Resistance to RXR-PNA in *rpsL*_*I82N*_ cells was not limited to a single target mRNA as the mutation also conferred resistance to a PNA directed against the AUG region of the *ftsZ* mRNA. RpsL_I82N_ mediated PNA resistance was also not limited to a single target within the mRNA translation initiation region as it applied to both PNA targeting the SD-sequence and those targeting the AUG region of *acpP* (Table 1).

Interestingly, the *rpsL*^I82N^ mutant displayed resistance to KFF-PNA, whereas the parental Evo-3 strain did not. Evo-3 carries insertion-sequence disruptions in *waaB* and *waaO*. Deletion of *waaB* and *waaO* increased sensitivity to KFF-PNA relative to the isogenic wild type (Table 1). Conversely, when *waaB* and *waaO* were deleted in the *rpsL*_*I82N*_ background, the MIC of KFF-PNA decreased to the wild-type level (Table 1). These results show that an intact LPS outer core contributes to intrinsic resistance to KFF-PNA in *E. coli* and that loss of the outer core masks the intracellular resistance conferred by *rpsL*_*I82N*_. The increased KFF-PNA susceptibility of the LPS mutants is consistent with increased entry of this conjugate, although uptake was not measured directly in these experiments.

### Several mutations in ribosomal protein S12 confer PNA resistance at a high fitness cost

To determine whether other *rpsL* mutations confer resistance to RXR-PNA, we combined a clinical-isolate screen with a focused genetic assessment. Approximately 2,300 clinical *E. coli* isolates were examined, and three isolates contained nonsynonymous *rpsL* variants (0.13%): *rpsL*_*T64A*_, *rpsL*_*I80L*_, and *rpsL*_*I82N*_. The isolate carrying *rpsL*_*I82N*_, recovered from a urinary-tract infection, was resistant to RXR-PNA, whereas the isolates carrying *rpsL*_*T64A*_ or *rpsL*_*I80L*_ were sensitive (Table 1). Because the clinical isolates were not isogenic, this comparison is supportive rather than causal; the resistance-conferring effect of *rpsL*_*I82N*_ is established by the isogenic reconstruction described above.

The *rpsL*_*I82N*_ mutant is positioned between two conserved regions of *rpsL* encoding residues 42-44 and 86-92 where mutations that confer resistance to streptomycin are commonly located [25], and in agreement *rpsL*_*I82N*_ cells are streptomycin sensitive (Table 1). Several substitutions affecting amino acids in or near position 82 of RpsL, including *rpsL*_*L74P*_, *rpsL*_*I80N*_, *rpsL*_*L81R*_, and *rpsL*_*I82F*_, increased RXR-PNA resistance relative to the wild type, with the strongest effect observed for *rpsL*_*I82F*_ (Table 1). The effect depended on the substituted side chain rather than position alone: at residue 80, I80L had no detectable effect, whereas I80N increased the MIC four-fold. Together, the clinical and targeted genetic data define a local region of S12, approximately residues 74-82, in which specific substitutions can promote PNA resistance.

Only *rpsL*_*I82N*_ was identified in the previous adaptive laboratory evolution experiment [8]. We hypothesized that this reflected simultaneous selection for PNA resistance and cellular fitness. Wild-type and Evo-3 cells had doubling times of 24 and 37 min in Mueller-Hinton broth II, respectively, demonstrating a substantial fitness cost in Evo-3. Expression of wild-type *rpsL* from pCA24n:*rpsL* reduced the Evo-3 doubling time from 37 to 24 min (Table 1), indicating that the main fitness cost in Evo-3 was associated with *rpsL*_*I82N*_. In agreement, cells carrying only *rpsL*_*I82N*_ had a doubling time of 33 min, whereas *rpsL*_*I82F*_ cells had a doubling time of 41 min. Thus, *rpsL*_*I82N*_ provided a favorable resistance-fitness trade-off among the tested alleles, which may explain its recovery during adaptive evolution.

### *rpsL*_*I82N*_ causes widespread changes in proteins subject to post-transcriptional control

To examine the effect of *rpsL*_*I82N*_ on global protein abundance, we compared the proteomes of the mutant and wild-type strains. In total, 2,073 of the 4,448 proteins in the *E. coli* UniProt reference proteome were identified with high confidence, across two replicates per strain. Of these, 1,097 proteins were significantly differentially abundant in the *rpsL*_*I82N*_ mutant using a false-discovery rate (FDR) below 0.5%. The abundance of 623 proteins increased, whereas 474 proteins decreased relative to the wild type.

A gene-regulator enrichment analysis indicated significant enrichment among proteins whose expression is regulated post-transcriptionally by base-pairing sRNAs or by ribosomal proteins (Table 2 and Supplementary Table S1). For example, GcvB has 26 annotated target proteins in the analysis, and all were altered in the *rpsL*_*I82N*_ dataset: 11 were significantly increased and 15 were significantly decreased in *rpsL*_*I82N*_ mutant cells (Table 2). A mixed pattern was also observed among RyhB-regulated proteins, whereas all quantified targets assigned to MicA and RydC regulons were increased (Table 2). We also observed that all proteins subjected to translational repression by the ribosomal proteins L1, S4, and S8 were significantly upregulated in the *rpsL*_*I82N*_ mutant cells relative to the wild type (Table 2).

**Table 2.**
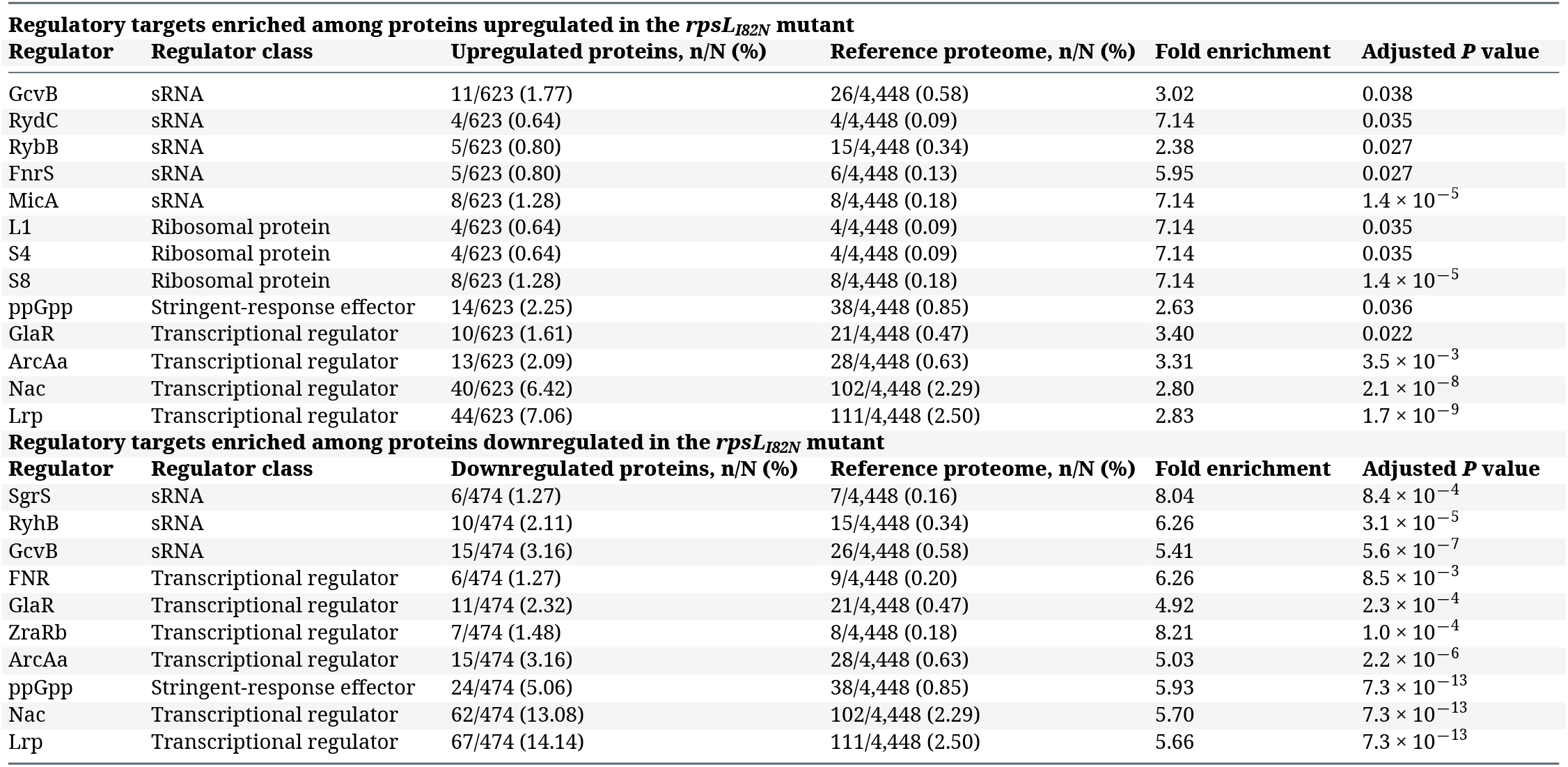
Regulatory-target enrichment was analysed separately among the 623 proteins significantly upregulated and the 474 proteins significantly downregulated in the *rpsL*_*I82N*_ mutant. The reference frequencies shown here were calculated using the 4,448 proteins in the *E. coli* reference proteome. Fold enrichment was calculated as the frequency in the differentially abundant protein set divided by the corresponding frequency in the reference proteome. Adjusted *P* values were corrected for multiple testing as described in Materials and Methods. sRNA, small regulatory RNA.

Together, the proteomic data indicates that the *rpsL*_*I82N*_ mutation is associated with broad changes in proteins under post-transcriptional control, including proteins regulated by base-pairing sRNAs and ribosomal proteins. Because the mutation substantially reduces growth rate and changes a large fraction of the quantified proteome, the enrichment analysis alone cannot distinguish direct effects on translation from indirect effects on transcription, mRNA stability, protein turnover, or growth-dependent proteome allocation.

### The effect of *rpsL*_*I82N*_ on sRNA-mediated regulation correlates with sRNA binding-site position

We next asked whether the effect of *rpsL*_*I82N*_ on sRNA-regulated protein abundance correlated with the location of the sRNA-binding site relative to the SD sequence and AUG start codon. To limit the analysis, we focused on proteins that were negatively regulated by an sRNA and differentially abundant in the *rpsL*_*I82N*_ mutant at an FDR below 0.1% (Supplementary Table S2).

Expression of proteins whose translation is negatively regulated by an sRNA with a binding site overlapping the SD sequence and/or AUG start codon was higher in the *rpsL*_*I82N*_ mutant relative to the wild type (Figure 1A). In contrast, proteins negatively regulated by an sRNA binding upstream or downstream of the SD/AUG region was lower in the mutant, with Fiu being the exception in this subset (Figure 1A). These observations suggested that the *rpsL*_*I82N*_ mutation affects how different sRNAs bind their respective targets, depending on the location of their binding sites on the mRNA.

**Figure 1.**
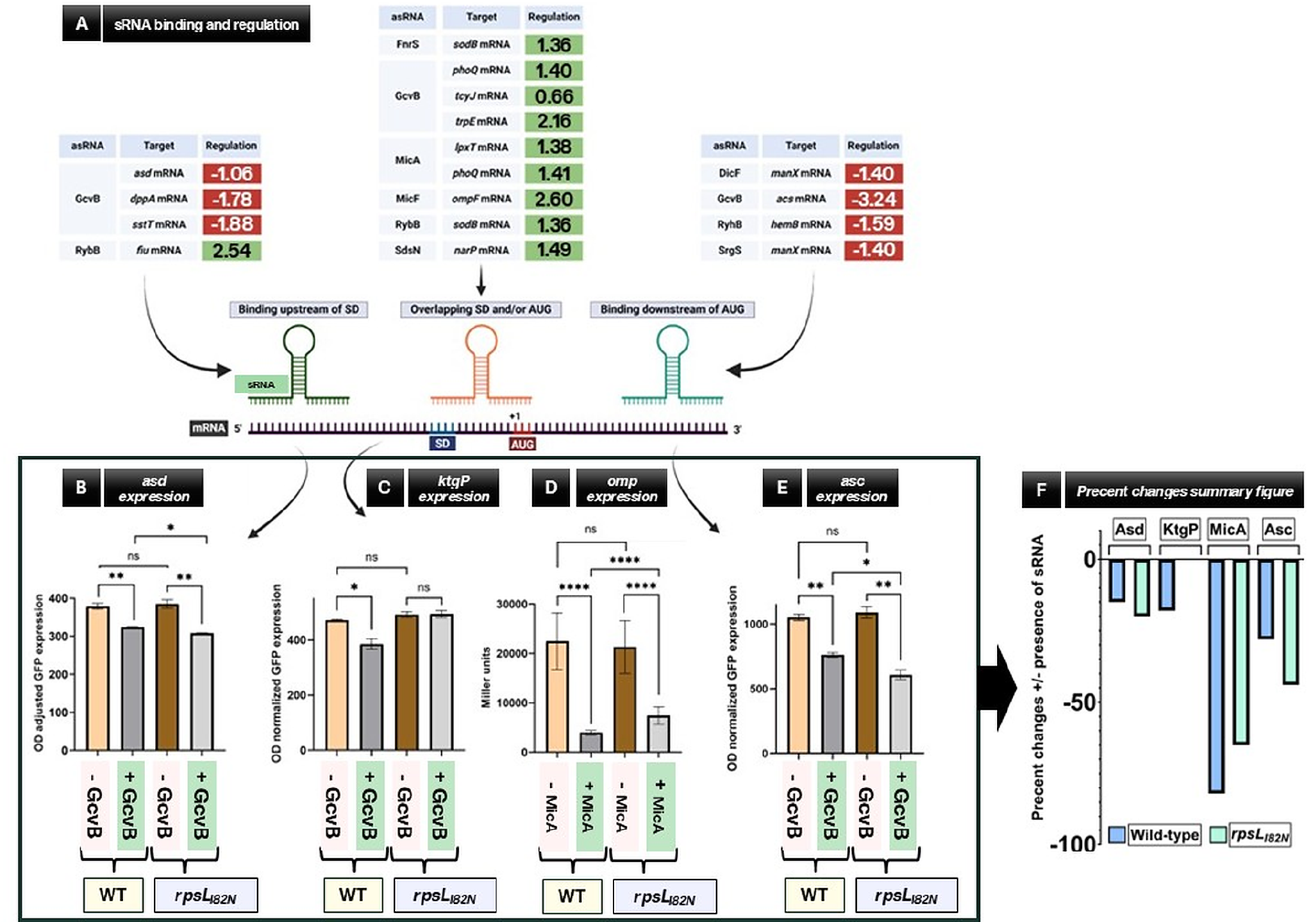
Effect of the *rpsL*_*I82N*_ mutation on sRNA-mediated regulation at different positions relative to the translation-initiation region. (A) Selected negatively regulated sRNA targets that were differentially abundant in the *rpsL*_*I82N*_ mutant relative to the wild type at a proteomic false-discovery rate below 0.1%. Targets were classified according to whether the reported sRNA-binding site was located upstream of the Shine–Dalgarno sequence, overlapped the Shine–Dalgarno sequence and/or AUG start codon, or was located downstream of the AUG. Values indicate the log_2_ fold change in protein abundance in the *rpsL*_*I82N*_ mutant relative to the wild type. Positive values are shown in green and negative values in red. The schematic indicates the relative positions of the sRNA-binding sites, Shine–Dalgarno sequence, AUG start codon and translational start position. (B) GcvB-dependent regulation of the pXG-10sf-*asd* translational reporter. Wild-type and *rpsL*_*I82N*_ cells carrying pXG-10sf-*asd* additionally contained either the vector control pTP11 or the GcvB-expression plasmid pPL-*gcvB*. GcvB reduced reporter expression in both the wild type (*P* = 0.0048) and *rpsL*_*I82N*_ mutant (*P* = 0.0056). Reporter expression did not differ significantly between the two strains in the presence of pTP11 (*P* = 0.6693), whereas expression in cells carrying pPL-*gcvB* was lower in the mutant than in the wild type (*P* = 0.0112). (C) GcvB-dependent regulation of the pXG-10sf-*kgtP* translational reporter. Wild-type and *rpsL*_*I82N*_ cells carrying pXG-10sf-*kgtP* additionally contained either pTP11 or pPL-*gcvB*. GcvB reduced reporter expression in the wild type (*P* = 0.0437), but no significant GcvB-dependent reduction was observed in the *rpsL*_*I82N*_ mutant (*P* = 0.9107). Basal reporter expression in the presence of pTP11 did not differ significantly between the two strains (*P* = 0.1902). (D) MicA-dependent regulation of the pOmpLac-M6 translational fusion. Wild-type and *rpsL*_*I82N*_ cells carrying pOmpLac-M6 additionally contained either the vector control pControl or the complementary MicA-expression plasmid pMicA-M6. MicA strongly reduced *ompA–lacZ* expression in the wild type (*P* < 0.0001) and in the *rpsL*_*I82N*_ mutant (*P* < 0.0001). Basal reporter expression did not differ significantly between the two strains carrying pControl (*P* = 0.5092), whereas reporter expression in the presence of pMicA-M6 was significantly higher in the *rpsL*_*I82N*_ mutant than in the wild type (*P* < 0.0001). (E) GcvB-dependent regulation of the pXG-10sf-*asc* translational reporter. Wild-type and *rpsL*_*I82N*_ cells carrying pXG-10sf-*asc* additionally contained either pTP11 or pPL-*gcvB*. GcvB reduced reporter expression in both the wild type (*P* = 0.0080) and the *rpsL*_*I82N*_ mutant (*P* = 0.0078). Basal reporter expression did not differ significantly between the two strains carrying pTP11 (*P* = 0.5550), whereas expression in cells carrying pPL-*gcvB* was lower in the mutant than in the wild type (*P* = 0.0167). (F) Relative change in reporter expression following sRNA expression. For each reporter and genotype, the percentage change was calculated relative to the corresponding vector-control condition as: Negative values indicate sRNA-mediated repression. Based on the group means shown in panels B–E, the changes in the wild type and *rpsL*_*I82N*_ mutant were, respectively, −14.5% and −20.0% for GcvB–*asd*, −18.4% and +0.4% for GcvB–*kgtP*, −82.4% and −65.2% for MicA–*ompA*, and −27.6% and −44.3% for GcvB–*asc*. Reporter fluorescence in panels B, C and E was normalized to OD_595_, whereas pOmpLac-M6 expression in panel D is reported as Miller units. Bars represent means and error bars represent SEM. Statistical comparisons were performed using two-tailed unpaired *t*-tests, with Welch’s correction where appropriate. Exact *P*-values are stated above. Significance symbols are defined as follows: ns, not significant; \**P* < 0.05; \*\**P* < 0.01; \*\*\**P* < 0.001; and \*\*\*\**P* < 0.0001.

To determine whether the proteomic pattern reflected altered sRNA regulation, we used plasmid-borne *lacZ* translational fusions for selected genes together with overproduction of the relevant sRNA from a co-resident plasmid. We selected *asd*, with a GcvB-binding site upstream of the SD/AUG region; *kgtP*, with a GcvB-binding site overlapping the SD/AUG region; *ompA*, with a MicA-binding site overlapping the SD/AUG region; and *asc*, with a GcvB-binding site downstream of the SD/AUG region. The use of three GcvB-regulated targets allowed the importance of binding-site position to be examined while keeping the regulatory sRNA constant.

In wild-type cells, overproduction of the relevant sRNA significantly reduced expression of *asd, kgtP, ompA*, and *asc* (Figures 1B-3D). This confirmed negative regulation by GcvB or MicA under the assay conditions. In *rpsL*_*I82N*_ cells, *asd, ompA*, and *asc* were also significantly downregulated following sRNA overproduction, whereas *kgtP* expression was unchanged in the presence or absence of GcvB (Figures 1B-3D).

The magnitude of repression differed between genes. The *asd* and *asc* translational fusions, whose GcvB-binding sites are located outside the SD/AUG region, were more strongly downregulated in *rpsL*_*I82N*_ than in wild-type cells (Figures 3B and 3E). Conversely, *kgtP*, whose GcvB-binding site overlaps the SD/AUG region, was not detectably downregulated by GcvB in the mutant (Figure 1C). Although *ompA* remained significantly repressed by MicA in the mutant, its expression in the presence of MicA was higher than in wild-type cells (Figure 1D).

Overall, the reporter data agreed with the position-dependent pattern observed in the proteomic analysis. The *rpsL*_*I82N*_ mutation reduced the ability of an sRNA to inhibit translation when the sRNA-binding site overlapped the SD/AUG region, but increased repression for the tested sites located outside this region (Figure 1F). This pattern also parallels the resistance to the anti-*acpP* and anti-*ftsZ* PNAs, which target the translation-initiation region.

#### Resistance to RXR-PNA does not correlate with classical decoding-fidelity phenotypes

Mutations in *rpsL* have traditionally been divided into hyper-accurate error-restrictive and error-prone ribosomal-ambiguity (ram) phenotypes. Error-restrictive alleles produce hyper-accurate ribosomes with increased rejection of near-cognate tRNAs, whereas ram alleles increase the rate of translational errors [14]. We used *lacZ* reporter constructs containing premature UGA (pSG3/4UGA) or UAG (pSG12-6) stop codons [26] to determine the translation-accuracy phenotype of *rpsL*_*I82N*_. The *rpsL*_*I82N*_ mutant produced significantly less β-galactosidase from both stop-codon reporters than the wild type (Figure 2A), consistent with an error-restrictive, hyper-accurate phenotype.

**Figure 2.**
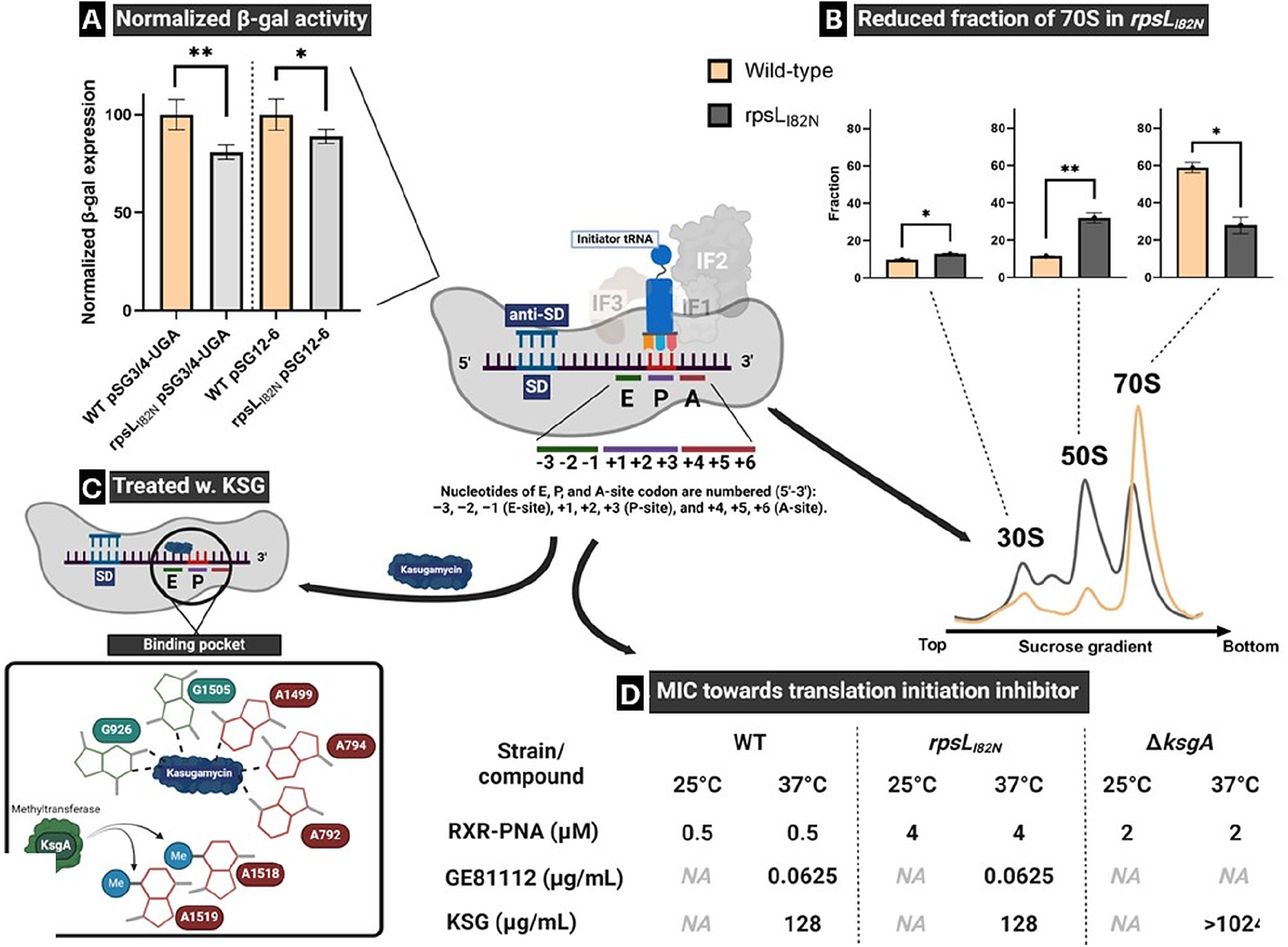
The *rpsL*_*I82N*_ mutation alters stop-codon readthrough and ribosome-state partitioning without conferring cross-resistance to kasugamycin or GE81112. (A) Classical decoding fidelity in wild-type and *rpsL*_*I82N*_ cells was assessed using the *lacZ* readthrough reporters pSG3/4UGA and pSG12-6, which contain premature UGA and UAG stop codons, respectively. β-Galactosidase activity was normalized to that of the corresponding wild-type strain, which was set to 100% for each reporter. The reduced reporter activity in the *rpsL*_*I82N*_ mutant indicates reduced stop-codon readthrough and is consistent with an error-restrictive, hyper-accurate phenotype. These assays do not assess initiator-tRNA selection or the kinetics of 30S initiation-complex formation. The central schematic illustrates the bacterial 30S translation-initiation complex, including the Shine–Dalgarno–anti-Shine–Dalgarno interaction, initiator tRNA, initiation factors IF1, IF2 and IF3, and the positions of the E-, P- and A-site codons relative to the AUG start codon. n = 4 for each condition. (B) Ribosome-state distribution in wild-type and *rpsL*_*I82N*_ cells determined by sucrose-density-gradient ultracentrifugation. A representative gradient profile is shown, with the positions of the 30S and 50S subunits and 70S ribosomes indicated. The bar graphs show quantification of the corresponding 30S, 50S and 70S fractions. Relative to the wild type, the *rpsL*_*I82N*_ mutant contained increased fractions of 30S and 50S particles and a reduced fraction of 70S ribosomes, consistent with altered subunit joining, 70S stability or ribosome assembly. An arrow indicates a small unassigned peak or shoulder between the 30S and 50S peaks in the mutant profile. Because this particle was not isolated or compositionally characterized, its identity remains unresolved; its presence is consistent with, but does not establish, defective ribosome assembly. n = 2 for each condition. (C) Schematic representation of the binding site of kasugamycin (KSG) within the 30S mRNA channel near the E- and P-site region. The enlarged schematic highlights nucleotides contributing to the kasugamycin-binding pocket and the adjacent KsgA-dependent dimethylation of A1518 and A1519 in 16S rRNA. The illustration is intended to show the relationship between kasugamycin-sensitive translation-initiation states and late-stage 30S maturation. (D) Minimum inhibitory concentrations of RXR-conjugated PNA, GE81112 and kasugamycin for wild-type, *rpsL*_*I82N*_ and Δ*ksgA* cells at 25°C and 37°C. The *rpsL*_*I82N*_ mutant showed an eightfold increase in the RXR–PNA MIC relative to the wild type at both temperatures. At 37°C, wild-type and *rpsL*_*I82N*_ cells displayed identical MICs for GE81112 and kasugamycin, indicating that the mutation did not confer detectable cross-resistance to these translation-initiation inhibitors. The Δ*ksgA* strain showed increased resistance to RXR– PNA and high-level resistance to kasugamycin. NA, not assessed. Bars represent mean ± SEM from independent biological experiments. Statistical comparisons were performed using two-tailed unpaired *t*-tests with Welch’s correction. \**P* < 0.05 and \*\**P* < 0.01.

The other PNA-resistant *rpsL* alleles were distributed between the two fidelity classes. *rpsL*_*L74P*_ and *rpsL*_*I82F*_ have error-restrictive phenotypes, whereas *rpsL*_*I80N*_ and *rpsL*_*L81R*_ have ram phenotypes [14]. Because PNA resistance occurred in both groups, increased RXR-PNA resistance did not correlate with the classical error-restrictive or ram phenotypes measured by stop-codon readthrough. These experiments do not exclude effects of the uS12 substitutions on translation-initiation fidelity or initiation-complex dynamics.

### *rpsL*_*I82N*_ shifts the ribosome profile toward 30S and 50S particles

Ribosomal protein S12 is located in the 30S subunit at the interface of functionally important 16S rRNA elements, and S12 has been implicated in RNA folding, small-subunit assembly and multiple stages of ribosome function [15,16,18,19]. We therefore examined whether *rpsL*_*I82N*_ altered the distribution of ribosomal particles.

Sucrose-gradient ultracentrifugation revealed a clear difference between the mutant and wild type. It showed a reduction in the 70S fraction (p < 0.015) and increases in the 30S and 50S fractions (p < 0.01 and p < 0.01, respectively; Figure 2B). In the representative *rpsL*_*I82N*_ profile, a small peak or shoulder was also visible between the assigned 30S and 50S peaks (arrow, Figure 2B). Because this particle was not isolated or compositionally characterized, it is referred to here as an unassigned intermediate particle; its presence is consistent with, but does not establish, defective ribosome assembly. Thus, *rpsL*_*I82N*_ alters ribosome-state partitioning and is consistent with impaired subunit joining, reduced 70S stability, altered ribosome biogenesis, or a combination of these effects. The gradient analysis does not distinguish mature free subunits from immature particles or mRNA-bound initiation complexes.

### Perturbation of 30S maturation and initiation increases PNA resistance

We next examined whether a second perturbation of 30S maturation influenced PNA susceptibility. KsgA dimethylates A1518 and A1519 in helix 45 of 16S rRNA and acts late in 30S assembly [20,21,27] (Figure 2C). Cells lacking KsgA accumulate altered or immature 30S particles, particularly at lower temperature. A Δ*ksgA* strain showed a four-fold increase in RXR-PNA resistance relative to the wild type at both 25°C and 37°C (Figure 2D). Published work has reported a larger free-30S fraction in Δ*ksgA* cells at 25°C than at 37°C, however the resistance to RXR-PNA did not change when tested at both temperatures. The PNA phenotype therefore did not scale directly with the bulk amount of 30S material and may depend on the structural or functional state of the particles.

Deletion of *ksgA* confers low-level resistance to kasugamycin, an antibiotic that perturbs mRNA positioning during translation initiation [13,28]. Treatment of wild-type cells with a sub-inhibitory concentration of kasugamycin increased RXR-PNA resistance four-fold, phenocopying the Δ*ksgA* strain (Figure 2D). This result is consistent with altered initiation-state dynamics influencing PNA activity, although a general effect of slower growth during sub-inhibitory antibiotic treatment cannot be excluded.

The *rpsL*_*I82N*_ mutant did not show increased resistance to kasugamycin or to GE81112, an initiation inhibitor that arrests a distinct 30S initiation intermediate (Figure 2D) [13].

## Discussion

We have shown that *rpsL*_*I82N*_ is a major intracellular determinant of resistance to antibacterial PNAs. The mutation increased resistance independently of the peptide carrier and protected against PNAs targeting different mRNAs and either the Shine–Dalgarno or AUG region. This adds intra-cellular PNA resistance through ribosomal changes to the previously described resistance originating from reduced entry [6–8]. Beyond synthetic PNAs, *rpsL*_*I82N*_ altered regulation by endogenous base-pairing sRNAs. The direction of this effect correlated with the location of the sRNA-binding site relative to the translation-initiation region. Regulation was weakened when the sRNA-binding site overlapped the SD/AUG region, but strengthened for the tested sites located outside this region. Finally, *rpsL*_*I82N*_ increased the 30S and 50S fractions and reduced the 70S fraction, while the representative mutant profile also contained a small unassigned intermediate particle. We therefore propose that altered uS12-dependent ribosome assembly and initiation-complex dynamics change access of both PNAs and sRNAs to their target sequences. Together, these findings identify ribosome occupancy of the translation-initiation region as a previously unrecognized determinant of susceptibility to synthetic and endogenous antisense regulators.

### *rpsL* defines a previously unrecognized intracellular resistance locus

The resistance phenotype was not unique to *rpsL*_*I82N*_. Substitutions at L74, I80, L81 and I82 increased RXR–PNA resistance, defining a local S12 region in which particular side-chain changes alter PNA susceptibility. The contrast between *rpsL*_*I80L*_ and *rpsL*_*I80N*_ is especially informative because it shows that the phenotype is not simply caused by changing any residue within this region. Rather, it depends on the physicochemical consequences of the individual substitution. The strongest resistance was observed with *rpsL*_*I82F*_, but this allele also imposed the largest measured growth defect. *rpsL*_*I82N*_ therefore appears to provide a favourable resistance–fitness compromise rather than the maximal resistance phenotype possible at this site.

This resistance–fitness relationship offers a plausible explanation for the preferential recovery of *rpsL*_*I82N*_ during adaptive evolution. Selection with increasing PNA concentrations would favour alleles that preserve sufficient synthesis of the PNA-targeted protein while retaining enough ribosomal capacity for continued growth. The 33-min doubling time of *rpsL*_*I82N*_ represents a substantial cost relative to the 24-min wild type, but the cost is lower than that of *rpsL*_*I82F*_. This is analogous to the general principle that resistance evolution selects on the combined phenotype of drug survival and growth rather than resistance magnitude alone. In the present system, the selected allele appears to occupy a local optimum in which PNA resistance is large enough to permit growth under selection but ribosome function is not as severely impaired as in higher-cost alternatives.

Identification of *rpsL*_*I82N*_ in a urinary-tract isolate shows that the allele can occur outside laboratory evolution. The clinical comparison itself is not causal because the three clinical isolates were not isogenic, but the reconstructed mutant establishes that the allele is sufficient to increase PNA resistance relative to an isogenic RpsL^+^ strain. The low frequency of nonsynonymous *rpsL* variants in the screened collection and the growth cost of *rpsL*_*I82N*_ suggest that these alleles may remain uncommon without a specific selective advantage. Nevertheless, the result is relevant to the future clinical development of PNA antibacterials because it demonstrates that naturally occurring ribosomal variation that alter PNA susceptibility exists even before therapeutic exposure.

### PNA resistance is distinct from the classical decoding-fidelity phenotype

The translation-fidelity experiments show that *rpsL*_*I82N*_ is an error-restrictive allele, but PNA resistance cannot be attributed simply to the classical decoding-fidelity phenotype measured by stop-codon readthrough. Other PNA-resistant S12 alleles were divided between error-restrictive and ribosomal-ambiguity phenotypes. This distinction is important because the best-known function of S12 mutations is their effect on tRNA selection and aminoglycoside susceptibility. Agarwal and colleagues similarly showed that S12 substitutions distributed throughout the protein can produce diverse fidelity phenotypes without necessarily conferring streptomycin resistance [14]. The present data therefore identify PNA susceptibility as an S12-dependent phenotype that is separable from the error-restrictive/ram classification. They do not, however, exclude effects on translation-initiation fidelity or initiation-complex dynamics.

This conclusion is consistent with growing evidence that uS12 influences RNA folding, ribosome assembly and translation initiation. Purified S12 can act as a broad-specificity RNA chaperone in vitro [15], and single-molecule analyses showed that S12 promotes cotranscriptional pre-16S rRNA folding and stable S4 recruitment during 30S assembly [16]. Moreover, uS12 substitutions at V32 and H76 evolved with an IF3 variant to optimize growth with an unconventional initiator tRNA, establishing genetic links between uS12, IF3 and initiator-tRNA selection [17]. H76 lies within the same local S12 region as several of the PNA-resistance substitutions identified here. Alteration of the S12 N-terminal extension can impair small-subunit assembly, and a mutation near the conserved PNSA loop has been shown to affect ribosome biogenesis, subunit association, initiation, elongation and recycling [18,19]. Although these substitutions are structurally distinct from I82N, they support a model in which *rpsL*_*I82N*_ may alter uS12-dependent rRNA folding and 30S assembly together with initiation-complex dynamics, rather than acting exclusively at the decoding step.

### Altered sRNA regulation links the PNA phenotype to target-site accessibility

The proteomic analysis revealed extensive physiological remodelling in the *rpsL*_*I82N*_ mutant. More than half of the quantified proteins showed significant changes, with enrichment of targets regulated by GcvB, MicA, RydC, RybB, RyhB and other sRNAs, as well as proteins subject to translational feedback by L1, S4 and S8. These results indicate that the mutation affects processes extending well beyond the two PNA-targeted transcripts. At the same time, the scale of the proteomic response means that enrichment alone cannot distinguish direct changes in translation from indirect consequences of slower growth, altered transcription, mRNA stability or protein turnover. The reporter experiments are therefore essential because they test defined sRNA–target interactions in the wild-type and mutant backgrounds.

The position-dependent pattern emerging from the proteome and reporters provides the most direct conceptual link between endogenous sRNA regulation and PNA resistance.

Direct competition between MicA and the 30S subunit for the *ompA* translation-initiation region has previously been demonstrated by toeprinting, providing a strong mechanistic precedent for the interpretation of the overlapping-site data [10]. The present work advances this principle by showing that competition can be altered by a mutation in the ribosome itself. In other words, the translational state of the target is not merely the process inhibited by an antisense molecule; it can also determine whether that molecule gains access to its binding site. This principle connects synthetic PNA action to endogenous post-transcriptional regulation and suggests that PNA resistance can emerge through the same kinetic competition that shapes natural sRNA activity.

The effect of sites outside the SD/AUG region is likely to be more mechanistically diverse. Bouvier and colleagues demonstrated that sRNA pairing within the early coding region can repress translation initiation even when the interaction does not directly occlude the canonical ribosome-binding site [11]. Other systems show that Hfq, RNA restructuring and recruitment of RNA-degradation machinery can allow sRNAs to act from a distance. Thus, “upstream,” “overlapping” and “downstream” should be viewed as useful functional classes rather than complete mechanistic descriptions. The relevant boundary may ultimately be whether the sRNA site is mutually exclusive with the complete 30S footprint or remains accessible in an mRNA-bound 30S complex. The Fiu exception and the mixed global response of GcvB targets also indicate that binding-site position is an important determinant but not the only one; RNA structure, SD strength, target abundance, Hfq dependence and decay pathways are also likely to influence the outcome. The reported RNA-chaperone activity of S12 also raises the possibility that *rpsL*_*I82N*_ changes local mRNA folding. However, the available studies examined purified S12 or pre-16S rRNA and do not establish direct remodelling of PNA- or sRNA-targeted mRNAs by ribosome-bound uS12 [15,16].

#### A ribosome-assembly and 30S-occupancy model for PNA resistance

The ribosome profiles provide a physical basis for a model in which altered ribosome assembly and initiation dynamics together influence PNA susceptibility. *rpsL*_*I82N*_ increased the 30S and 50S fractions while reducing the 70S fraction, and the representative mutant profile contained a small unassigned particle between the 30S and 50S peaks. This distribution is consistent with impaired or delayed subunit joining, reduced stability of 70S ribosomes, altered ribosome biogenesis or a combination of these factors. The position of the unassigned particle alone does not establish whether it is an immature 30S or 50S particle, another ribonucleoprotein complex or an experimental feature. Likewise, the gradients do not establish that the increased 30S peak consists of mature initiation-competent subunits, nor do they distinguish free mature subunits from immature particles or mRNA-bound initiation complexes. Nevertheless, they demonstrate that the mutation changes the abundance and heterogeneity of ribosomal particles and their partitioning into 70S ribosomes.

We propose that *rpsL*_*I82N*_ first alters uS12-dependent rRNA folding and/or 30S assembly, producing an altered and potentially heterogeneous population of ribosomal particles. Within the subset of initiation-competent 30S particles, one or more of the following changes may increase effective occupancy of transiently exposed translation-initiation regions (Figure 3):

**Figure 3.**
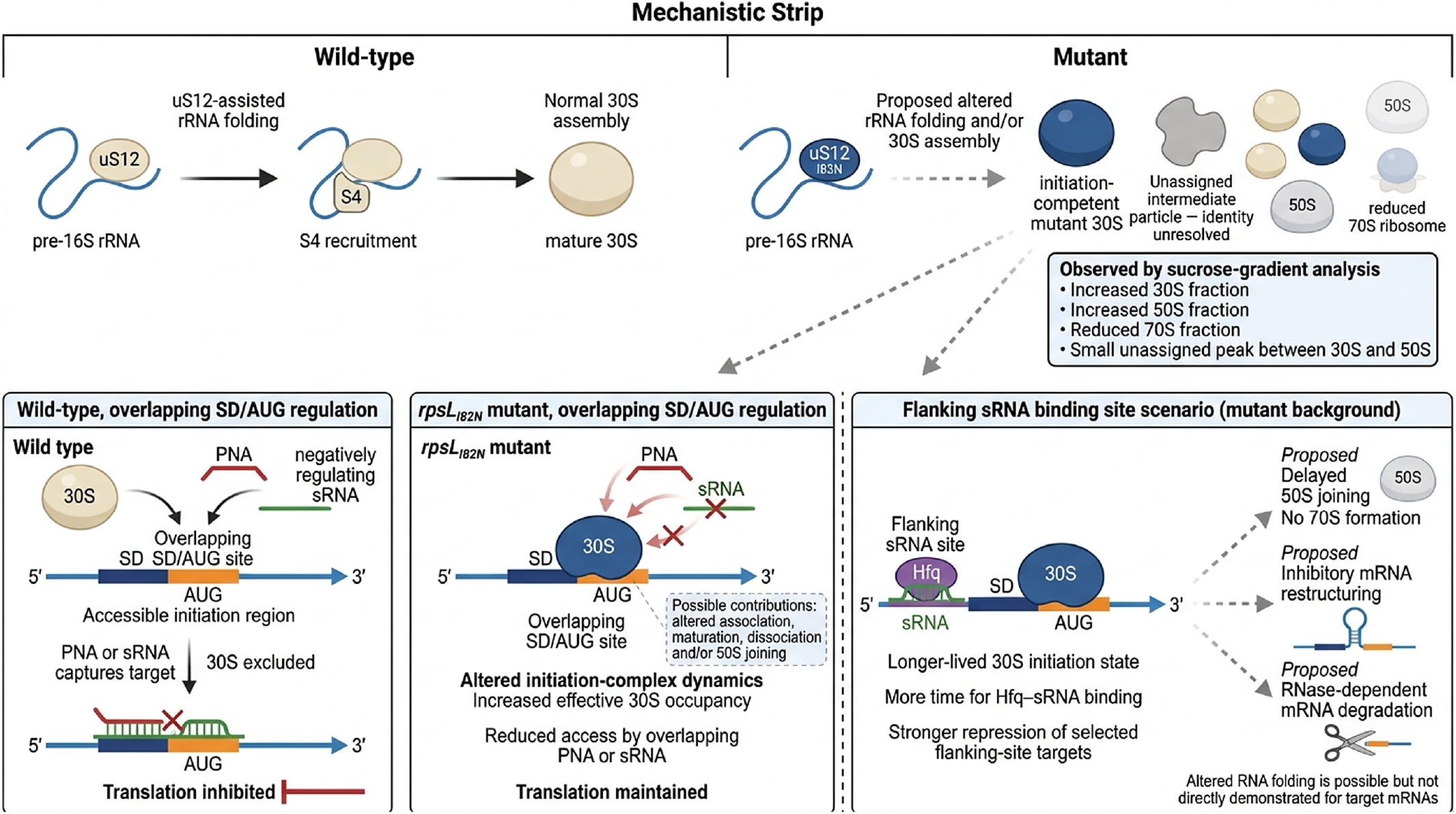
Proposed model linking altered uS12-dependent ribosome assembly and 30S initiation-state dynamics to position-dependent PNA and sRNA regulation. The *rpsL*_*I82N*_ substitution is proposed to alter S12-dependent rRNA folding and 30S assembly, generating an altered and potentially heterogeneous population of ribosomal particles. The sucrose-gradient profile supports altered particle partitioning but does not establish the identity or translation competence of the accumulated 30S material or the unassigned intermediate particle. In wild-type cells, the translation-initiation region remains accessible for sufficient time to allow a PNA or negatively regulating sRNA whose binding site overlaps the Shine–Dalgarno sequence and/or AUG start codon to hybridize to the target mRNA. Antisense binding excludes the 30S subunit and inhibits translation. In the *rpsL*_*I82N*_ mutant, altered association, maturation, dissociation and/or 50S-joining kinetics of initiation-competent 30S particles are proposed to increase effective occupancy of the SD/AUG region. This reduces access by PNAs and sRNAs targeting overlapping sequences and permits continued translation. The dashed vertical line separates this overlapping-site scenario from the proposed effect on sRNAs binding outside the 30S footprint. When an Hfq–sRNA complex targets a flanking region upstream or downstream of the SD/AUG site, a longer-lived 30S initiation state may increase the time available for sRNA binding and strengthen repression by delaying 50S joining, promoting inhibitory mRNA restructuring or facilitating RNase-dependent mRNA degradation. S12 RNA-chaperone activity also raises the possibility of altered RNA folding, but direct remodelling of target mRNAs by ribosome-bound uS12 has not been demonstrated. Red crosses indicate inhibited interactions or transitions, whereas dashed arrows indicate proposed or indirect effects. PNAs and sRNAs in the overlapping-site panels represent alternative antisense regulators rather than simultaneous binding events.

- a larger effective pool of initiation-competent 30S particles available for mRNA capture;
- faster association of 30S with the target transcript;
- slower dissociation of the 30S complex;
- delayed maturation of the 30S initiation complex;
- or slower 50S joining, which prolongs the lifetime of a pre-70S state.

The current data do not distinguish among these possibilities and do not establish that the larger total 30S fraction itself causes resistance. SD recognition is mediated by pairing with the anti-Shine–Dalgarno sequence of 16S rRNA. The more defensible interpretation is therefore that *rpsL*_*I82N*_ alters the effective occupancy time or capture probability of the translation-initiation region.

For PNAs and sRNAs whose binding sites overlap the occupied 30S footprint, ribosome binding and antisense binding are mutually exclusive. Increased 30S occupancy would narrow the temporal window available for antisense hybridization, allowing a larger fraction of transcripts to remain translationally competent. The mutant ribosome is unlikely to displace a fully formed high-affinity PNA–RNA duplex. Rather, resistance would arise because the ribosome captures the site before the PNA binds. This model predicts a strong order-of-addition effect in a purified system: *rpsL*_*I82N*_ 30S particles should protect the target more effectively when allowed to bind before or simultaneously with PNA, but should provide little advantage when PNA is prebound.

For an sRNA site that remains accessible while the 30S subunit occupies the translation-initiation region, prolonged residence of a 30S intermediate could have the opposite consequence. A longer-lived intermediate would provide more time for the sRNA–Hfq complex to bind an upstream or downstream site, alter the local mRNA structure to facilitate sRNA binding or both. The resulting complex might interfere with a conformational transition required for productive initiation, delay 50S joining, restructure the mRNA or promote RNase-dependent degradation. This arm of the model is more speculative than the direct-competition mechanism and should be presented as a working hypothesis. It nevertheless provides a coherent explanation for why the same ribosomal mutation can protect overlapping sites while strengthening repression at selected flanking sites.

### KsgA and initiation inhibitors support convergence at the 30S initiation pathway

The Δ*ksgA* phenotype independently connects PNA susceptibility to small-subunit maturation. KsgA acts late in 30S biogenesis, and structural studies show that KsgA binding prevents helix 44 from adopting its mature position until late assembly steps have been completed [20,21]. The four-fold PNA resistance of Δ*ksgA* is therefore consistent with the conclusion that the structural and functional state of the 30S population influences PNA activity. The identical increase in resistance at 25°C and 37°C, despite reported temperature-dependent changes in the abundance of immature particles, argues against a simple model in which resistance scales directly with the total 30S peak. The qualitative state of the particles appears more important than bulk subunit abundance.

Sub-inhibitory kasugamycin also increased PNA resistance, but this observation should not be taken to mean that kasugamycin, Δ*ksgA* and *rpsL*_*I82N*_ generate an identical ribosomal structure. Recent cryo-EM analysis shows that kasugamycin and GE81112 act at distinct stages of 30S initiation-complex formation: kasugamycin perturbs an early stage of stable initiator-tRNA accommodation and start-codon recognition, whereas GE81112 permits AUG recognition but prevents efficient progression towards 70S initiation-complex formation [13]. The absence of cross-resistance to either compound in *rpsL*_*I82N*_ indicates that the mutation does not simply prevent drug binding or create a constitutively initiation-inactive ribosome. Instead, several distinct perturbations of 30S maturation or initiation-state distribution appear capable of reducing PNA susceptibility.

#### Relevance to antibacterial PNA resistance

Whereas Cpx activation, SbmA loss and LPS alterations affect how much active PNA reaches the cytoplasm, the *rpsL*_*I82N*_ mutation alters ribosome–mRNA interactions at the translation-initiation region after PNA entry. This provides a possible explanation for why whole-cell PNA activity cannot always be predicted from target abundance or duplex stability alone.

These findings have practical implications for PNA development. Screening pipelines currently emphasize sequence complementarity, predicted duplex stability, target essentiality and delivery. The present results indicate that target-site accessibility should also be considered in the context of ribosome occupancy, initiation strength and bacterial physiological state. Testing multiple target positions, measuring direct target-protein knockdown and monitoring intracellular PNA accumulation will help distinguish poor uptake from poor target engagement. Combining PNAs that act through different transcripts or target-site geometries may also reduce the probability that a single ribosomal alteration protects all components of a treatment.

In conclusion, *rpsL*_*I82N*_ defines an intracellular route to antibacterial PNA resistance that operates at the ribosome– mRNA interface. The mutation changes ribosome-state partitioning, alters position-dependent sRNA regulation and protects multiple translation-initiation-region targets independently of peptide carrier. The data establish bacterial PNA susceptibility as the product of sequential envelope and intracellular barriers. More broadly, they identify translation-initiation-site occupancy as a determinant of whether synthetic and endogenous antisense regulators can gain access to bacterial mRNA.

## Methods

### Growth conditions

Cells were grown in Luria-Bertani (LB) medium or Müller-Hinton II broth (MHBII) at 37 °C with aeration unless otherwise stated. When required, antibiotics were added at the following concentrations: chloramphenicol, 20 µg/mL; ampicillin, 150 µg/mL; kanamycin, 50 µg/mL; streptomycin, 100 µg/mL; and tetracycline, 10 µg/mL [8].

### Bacterial strains and plasmids

All bacterial strains and plasmids used in this study are listed in Supplementary Tables S3 and S4, respectively. PNA compounds are listed in Supplementary Table S5, and oligonucleotide sequences are listed in Supplementary Table S6. Deletion alleles and overexpression plasmids were obtained from the Keio collection [29] and ASKA library [30], respectively. pCA24n-based plasmids were transformed into *E. coli* MG1655 with selection for chloramphenicol resistance, and expression was induced with 0.1 mM isopropyl β-D-1-thiogalactopyranoside (IPTG).

#### Construction of chromosomal *rpsL* mutants

The *rpsL*_*I82N*_ and *rpsL*_*I82F*_ substitutions were introduced into MG1655 using the two-plasmid CRISPR-Cas9/λ-Red system described by Jiang et al. [31]. The *rpsL*-specific guide sequence was introduced into pTarget using primers pTarget_sgRNA_*rpsL*mut_FW and pTarget_sgRNA_RV (Supplementary Table S6), generating pTarget:*rpsL* (pJFM40). The plasmid was verified by Sanger sequencing. Single-stranded donor oligonucleotides Oligo_I82N and Oligo_I82F were used to introduce the respective substitutions.

MG1655 carrying pCas was induced with 0.2% arabinose during preparation of electrocompetent cells. pJFM40 (100 ng) and 400 ng of the relevant donor oligonucleotide were mixed with the electrocompetent cells and electroporated in a 0.1-cm cuvette at 1.8 kV. Cells were recovered in 1 mL LB for 1 h at 32 °C and plated on LB agar containing kanamycin and streptomycin. Candidate colonies were screened by PCR and Sanger sequencing. pJFM40 was subsequently counter-selected, and the temperature-sensitive pCas plasmid was removed by growth at 37 °C without selection. This generated MG1655 *rpsL*_*I82N*_ and MG1655 *rpsL*_*I82F*_. Both alleles were confirmed by Sanger sequencing.

To combine the *cpxR*_*L20Q*_ and *rpsL*_*I82N*_ substitutions, *rpsL*_*I82N*_ was introduced into MG1655 *cpxR*_*L20Q*_ using the same CRISPR-Cas9/λ-Red procedure [8,31]. The *waaB*/*waaO* deletion was introduced into the MG1655 *rpsL*_*I82N*_ background by λ-Red recombination as described previously [8,32], generating MG1655 *rpsL*_*I82N*_ Δ*waaBO*.

#### Transfer of additional S12 alleles and construction of Δ*ksgA*

MC41 derivatives carrying *rpsL*_*K44I*_, *rpsL*_*L74P*_, *rpsL*_*S78C*_, *rpsL*_*V79E*_, *rpsL*_*I80N*_, *rpsL*_*L81R*_, *rpsL*_*I82F*_, *rpsL*_*R86H*_, and *rpsL*_*R86S*_ were obtained from Michael O’Connor [14,33]. A Δ*aroE* allele obtained from the Keio collection [29] was transferred into MG1655 by P1 transduction [34]. Because *aroE* is closely linked to *rpsL* and is required for growth on glucose minimal medium, P1 lysates prepared from the MC41 donor strains were used to transduce MG1655 Δ*aroE*; transductants that had reacquired *aroE* were selected on glucose minimal medium. The transferred *rpsL* alleles were confirmed by Sanger sequencing. The MG1655 *rpsL*_*I82F*_ strain used in the present study was generated by CRISPR-Cas9 as described above.

For the kasugamycin experiments, the Δ*ksgA* allele was obtained from the Keio collection [29] and transferred into MG1655 by P1 transduction [34]. The resulting MG1655 Δ*ksgA* strain was verified by PCR.

#### Construction of low-copy-number *rpsL* plasmids

The chromosomal *rpsL* region was PCR-amplified from MG1655 and MG1655 *rpsL*_*I82N*_ using primers *rpsL*_pALO277_FW and *rpsL*_pALO277_RV (Supplementary Table S6). The resulting amplicons were cloned into the low-copy F-based plasmid pALO277 [24] by restriction-ligation cloning, generating pALO277:*rpsL* (pJFM41) and pALO277:*rpsL*_*I82N*_ (pJFM42), respectively. Both plasmids were verified by Sanger sequencing. pALO277 is maintained at approximately one to two copies per genome equivalent [23,24].

### PNA-peptide conjugates and antimicrobial compounds

PNA-peptide conjugates were synthesised as described previously [1,7,8]. Peptide amino acids are shown in upper-case letters and PNA nucleobases in lowercase letters. Ahx denotes 6-aminohexanoic acid and eg1 denotes 8-amino-3,6-dioxaoctanoic acid. The anti-*acpP* RXR-PNA was H-(R-Ahx-R)_4_-Ahx-(βAla)-ctcatactct-NH_2_; the anti-*acpP* KFF-PNA was H-(KFF)_3_K-eg1-ctcatactct-NH_2_; and the anti-*ftsZ* RXR-PNA was H-(R-Ahx-R)_4_-Ahx-(βAla)-ttcaaacatagt-NH_2_ [8]. Naked anti-*acpP* PNA contained the sequence H-ctcatactct-NH_2_. The compounds used in this study are listed in Supplementary Table S5.

PNA stocks were dissolved in 0.02% acetic acid containing 0.4% bovine serum albumin (BSA), and twofold working dilutions were prepared in 0.01% acetic acid containing 0.2% BSA, as described previously [8]. Low-binding micro-centrifuge tubes, low-retention pipette tips and low-binding microtitre plates were used throughout PNA handling. Final stock concentrations were determined by absorbance at 260 nm.

Kasugamycin was obtained from Sigma-Aldrich, and GE81112 was obtained from Fisher Scientific. The compounds were prepared according to the manufacturers’ instructions.

### Determination of minimum inhibitory concentrations

Minimum inhibitory concentrations (MICs) were determined by broth microdilution using a modification of standard procedures [8,35,36]. Briefly, an overnight bacterial culture was diluted to approximately 5 × 10^5^ CFU/mL in MHBI. Bacterial suspension (100 µL) was dispensed into a low-binding 96-well microtitre plate (Thermo Scientific, catalogue no. 260895) together with 11 µL of test compound prepared as a twofold serial dilution. Vehicle-control wells contained 100 µL bacterial suspension and 11 µL PNA diluent. Plates were incubated without shaking for 18–24 h at 37 °C unless otherwise stated. The MIC was defined as the lowest concentration that prevented visible growth, corresponding to OD_595_ < 0.1. Inoculum density was verified by viable counting in representative experiments.

MHBI was supplemented with 0.1 mM IPTG when strains carried pCA24n-based plasmids. Naked PNA was tested in the AS19 background [22] using the same endpoint. Temperature-dependent MICs for the wild type, *rpsL*_*I82N*_ and Δ*ksgA* were determined at 25 °C and 37 °C. For the kasugamycin-perturbation experiment, wild-type cells were assayed for RXR-PNA susceptibility in the continuous presence of a sub-inhibitory concentration of kasugamycin.

### Growth measurements and doubling-time calculation

Doubling times were determined from three independent cultures grown in MHB II at 37 °C with aeration. Overnight culture (10 µL) was inoculated into 50 mL medium in a 250-mL flask. OD_595_ was measured during exponential growth, between approximately OD 0.05 and 0.5, at 5–10-min intervals. The specific growth rate (µ) was obtained from the slope of ln(OD) versus time over the exponential phase, and the doubling time was calculated as ln(2)/µ. Cultures carrying pCA24n:*rpsL* contained 0.1 mM IPTG and the appropriate selective antibiotic.

### Screening of clinical *Escherichia coli* isolates

Approximately 2,300 clinical *Escherichia coli* isolates were screened for nonsynonymous variation in *rpsL*. Isolates carrying candidate alleles were tested for RXR-PNA susceptibility using the MIC procedure described above.

### β-Galactosidase activity and Miller-unit calculation

β-Galactosidase activity was measured using O-nitrophenyl-β-D-galactopyranoside (ONPG) and expressed as Miller units [34]. Overnight culture (4 µL) was inoculated into 20 mL LB in a 100-mL flask and incubated at 37 °C with aeration. Beginning at approximately OD_600_ 0.1, five samples were collected on ice at 5–10-min intervals. Cells were permeabilised with toluene. An appropriate volume of permeabilised culture was added to 1 mL of 0.8 mg/mL ONPG in Z-buffer containing 60 mM Na_2_HPO_4_·7H_2_O, 40 mM NaH_2_PO_4_·H_2_O, 100 mM KCl, 1 mM MgSO_4_ and 50 mM β-mercaptoethanol, adjusted to Ph 7.0. Reactions were incubated at 30 °C and stopped with 0.5 mL 1 M Na_2_CO_3_ after development of a yellow colour. Samples were clarified by centrifugation at 15,000 × g for 5 min, and OD_420_ was measured.

Miller units were calculated as 1,000 × OD_420_/(t × V × OD_600_), where t is the reaction time in minutes and V is the volume of permeabilised culture in millilitres. No OD_550_ correction was applied.

#### Translation-fidelity reporters

Stop-codon readthrough was assessed using pSG3/4UGA and pSG12-6, which contain premature UGA and UAG codons within *lacZ*, respectively [26]. The reporter plasmids were introduced into the wild-type and *rpsL*_*I82N*_ backgrounds, and β-galactosidase activity was determined as described above. Activity was normalised to the corresponding wild-type mean, which was set to 100% for each reporter. Four independent biological experiments were analysed for each condition.

#### MicA-dependent regulation of the *ompA*-*lacZ* reporter

MicA-dependent regulation was analysed using the matched M6 reporter pair described by Udekwu et al. [10]. Wild-type and *rpsL*_*I82N*_ cells carried pOmpLac-M6 together with either pControl or pMicA-M6. pOmpLac-M6 contains a six-base substitution in the MicA target sequence, and pMicA-M6 contains the complementary substitutions. β-Galactosidase activity was determined using the ONPG assay and expressed as Miller units. The experiment was performed on three independent days. Measurements obtained within each day were averaged before statistical analysis, and experimental day was treated as the biological replicate and blocking factor.

### GcvB-dependent sfGFP translational reporters

GcvB regulation of *asd, kgtP* and *asc* was analysed using pXG-10sf translational fusions together with the vector control pTP11 or the GcvB-expression plasmid pPL-*gcvB*, based on the system described by Miyakoshi et al. [12]. Reporter plasmids were introduced into wild-type and *rpsL*_*I82N*_ cells. Fluorescence (excitation, 485 nm; emission, 535 nm) and OD_595_ were recorded at 37 °C in a microplate reader. For each well and measurement cycle, fluorescence was divided by the corresponding OD_595_. Technical wells were averaged within each independent experiment before statistical analysis.

### Sample preparation for LC-MS/MS

Quantitative proteome analysis was performed using data dependent acquisition liquid chromatography tandem mass spectrometry (DDA LC-MS/MS). Two independent biological cultures of wild-type and *rpsL*_*I82N*_ cells were grown under identical conditions and harvested during exponential growth at OD_595_ 0.2. Cell pellets were washed in 0.9% saline buffer, frozen and stored at −80 °C. Cells were lysed in 100 ul 8 M urea – 100 mM ammonium bicarbonate followed by sonication in a Bioruptor Plus (Diagenode) sonicator at 4 °C for 40 cycles with 15 sec on, 15 sec off. Insoluble material was removed by centrifugation at 13 000 rpm for 10 min at 4 °C. Disulfide bonds were reduced with 5 mM tris(2-carboxyethyl)phosphine for 30 min at 37 °C, and cysteine residues were alkylated with 10 mM iodoacetamide for 60 min at room temperature in the dark. Samples were diluted with 100 mM ammonium bicarbonate to a final urea concentration of 1.5 M and digested with sequencing-grade trypsin (Promega, V5111) for 18 h at 37 °C. Digestion was terminated by acidification with formic acid to a final pH 3. Peptides were purified using C18 reversed-phase columns, dried under vacuum and reconstituted in 0.2% formic acid, 2% acetonitrile prior to LC-MS/MS analysis.

#### Liquid chromatography tandem mass spectrometry (LC–MS/MS)

All peptides were analyzed on an Eclipse mass spectrometer connected to an ultra-high performance Ultimate3000 liquid chromatography system (both Thermo Scientific). Approximately 500 ng of peptides were separated on a Thermo EASY-Spray column (Thermo Scientific 25 cm column, column temperature 45 °C) operated at a maximum pressure of 900 bar. A non-linear gradient of 80% acetonitrile in aqueous 0.1% formic acid was run for 120 min at a flow rate of 300 nl/min. One full MS scan (resolution 120,000) for a mass range of 350-1400 m/z was followed by MS/MS scans resolution 15,000 m/z. The cycle time was 3 sec. The precursor ions were isolated with 1.6 m/z isolation window and fragmented using higher-energy collisional-induced dissociation (HCD) at a normalized collision energy of 30. The dynamic exclusion was set to 45 or 60 sec.

#### Proteomic data processing and statistical analysis

Raw LC-MS/MS data were processed using Proteome Discoverer 2.5 and searched against the *Escherichia coli* reference proteome (Uniprot ID UP000000625) downloaded from UniProt in March 2022, supplemented with common contaminant sequences and reverse decoy sequences. Trypsin specificity was specified, allowing up to 2 missed cleavages. Carbamidomethylation of cysteine residues was defined as a fixed modification, and oxidation of methionine residues as a variable modification. Peptide-spectrum matches and protein identifications were controlled at a false-discovery rate of 1% at the peptide and protein levels.

Abundance values were Log2 transformed and mean normalized, and differential abundance between the *rpsL*_*I82N*_ mutant and wild type was assessed using Perseus.

### Regulator enrichment and sRNA-binding-site classification

**Regulator-target enrichment was performed separately among proteins increased and decreased in the *rpsL***_***I82N***_ **mutant at proteomic FDR < 0.5%. Regulatory annotations were obtained from EcoCyc (v26.0). Swiss-Prot identifiers were mapped via the association table to EcoCyc entities, with transcription-unit definitions expanded to the gene level to retain regulatory interactions targeting the gene or its transcription unit, thereby enabling access to the regulator identity, mode (activation/repression), and interaction type according to EcoCyc ontology. Regulator overrepresentation among the up- and down-regulated subsets was tested against a background of 4 392 *E. coli* protein-coding genes using a one-tailed hypergeometric test, followed by Bonferroni correction, in Python. For the positional analysis, negatively regulated sRNA targets that were differentially abundant at FDR < 0.1% were classified as binding upstream of the SD/AUG region, overlapping the SD and/or AUG, or downstream of the AUG**.

### Sucrose-gradient analysis of ribosomal particles

Ribosomal particles were analysed by sucrose-density-gradient ultracentrifugation using a procedure supplied by Thomas Prossliner [37]. Wild-type and *rpsL*_*I82N*_ cultures were grown to mid-exponential phase, rapidly chilled and harvested by centrifugation at 4 °C. Cell pellets were resuspended in ice-cold ribosome lysis buffer containing 25 mM HEPES (pH 7.5), 100 mM KOAc, 15 mM Mg(OAc)_2_ and 1 mM

DTT. An equal volume of zirconia/silica beads (BioSpec) was added, and lysis was performed by five cycles of 1 min vortexing followed by 1 min of cooling on ice. Cellular debris was removed by centrifugation at 12000×g for 5 min at 4 °C. Clarified lysate corresponding to 20 A260 units was diluted to a volume of 300 µl in lysis buffer and layered onto continuous 5-40% (w/v) sucrose gradients prepared in lysis buffer supplemented with 0.01% n-dodecyl-d-maltoside.

Gradients were centrifuged at 41000 rpm/287660×g for 3 h at 4 °C in a TH-641 rotor using a Thermo Sorvall WX90 ultracentrifuge. Following centrifugation gradients were fractionated using a Gradient Fractionator (BioComp) and ribosomal particles were detected by continuous monitoring at 254 nm. Peaks corresponding to free 30S and 50S subunits and 70S ribosomes were assigned from their sedimentation positions. Profiles were background-corrected, and peak areas were normalised to the total integrated absorbance signal before comparison between strains. Two independent biological experiments were analysed.

### Statistical analysis

Statistical analyses were performed using GraphPad Prism 9.0 (GraphPad Software, San Diego, CA, USA). Independent biological experiments were treated as the experimental unit, and technical measurements were averaged within each biological experiment before analysis unless a hierarchical or repeated-measures model was used. Pairwise comparisons were performed using two-tailed tests with Welch’s correction where appropriate. The exact test, biological replicate number and definition of error bars are stated in the corresponding figure legends. Differences were considered significant at *P* < 0.05 after the stated multiple-comparison correction.

## Supplementary reporting tables

**Supplementary Table S3.** Bacterial strains.

| Relevant genotype | Abbreviated form | Source/reference |
| --- | --- | --- |
| MG1655; F<sup>−</sup> λ<sup>−</sup> rph-1 | Wild type | [38] |
| MG1655 evolved during RXR-PNA selection; *cpxR*<sub>L20Q</sub>, *rpsL*<sub>I82N</sub>, IS1::*waaB*, IS1::*waaO* | Evo-3 | [8] |
| *cpxR*<sub>L20Q</sub><sup>a</sup> | *cpxR*<sub>L20Q</sub> | [8] |
| *rpsL*<sub>I82N</sub><sup>a</sup> | *rpsL*<sub>I82N</sub> | This study |
| *cpxR*<sub>L20Q</sub> *rpsL*<sub>I82N</sub><sup>a</sup> | *cpxR*<sub>L20Q</sub> *rpsL*<sub>I82N</sub> | This study |
| Δ*waaBO*<sup>a</sup> | Δ*waaBO* | [8] |
| *rpsL*<sub>I82N</sub> Δ*waaBO*<sup>a</sup> | *rpsL*<sub>I82N</sub> Δ*waaBO* | This study |
| AS19 | AS19 | [22] |
| Δ*aroE*<sup>a</sup> | Δ*aroE* | This study |
| MC41 *rpsL*<sub>K44I</sub> |  | [14] |
| MC41 *rpsL*<sub>L74P</sub> |  | [14,33] |
| MC41 *rpsL*<sub>S78C</sub> |  | [14] |
| MC41 *rpsL*<sub>V79E</sub> |  | [14] |
| MC41 *rpsL*<sub>I80N</sub> |  | [14] |
| MC41 *rpsL*<sub>L81R</sub> |  | [14] |
| MC41 *rpsL*<sub>I82F</sub> |  | [14] |
| MC41 *rpsL*<sub>R86H</sub> |  | [14] |
| MC41 *rpsL*<sub>R86S</sub> |  | [14] |
| *rpsL*<sub>K44I</sub><sup>a</sup> | *rpsL*<sub>K44I</sub> | This study |
| *rpsL*<sub>L74P</sub><sup>a</sup> | *rpsL*<sub>L74P</sub> | This study |
| *rpsL*<sub>S78C</sub><sup>a</sup> | *rpsL*<sub>S78C</sub> | This study |
| *rpsL*<sub>V79E</sub><sup>a</sup> | *rpsL*<sub>V79E</sub> | This study |
| *rpsL*<sub>I80N</sub><sup>a</sup> | *rpsL*<sub>I80N</sub> | This study |
| *rpsL*<sub>L81R</sub><sup>a</sup> | *rpsL*<sub>L81R</sub> | This study |
| *rpsL*<sub>I82F</sub><sup>a</sup> | *rpsL*<sub>I82F</sub> | This study |
| *rpsL*<sub>R86H</sub><sup>a</sup> | *rpsL*<sub>R86H</sub> | This study |
| *rpsL*<sub>R86S</sub><sup>a</sup> | *rpsL*<sub>R86S</sub> | This study |
| Δ*ksgA*<sup>a</sup> | Δ*ksgA* | This study; Keio donor [29] |

<sup>a</sup> Otherwise as wild-type

**Supplementary Table S4.** Plasmids.

| **Plasmid** | **Relevant feature** | **Source/reference** |
| --- | --- | --- |
| pCA24n | ASKA vector; pT5-lac promoter | [30] |
| pCA24n:*rpsL* | IPTG-inducible wild-type *rpsL* | [30] |
| pALO277 | Low-copy F-based plasmid | [23,24] |
| pALO277:*rpsL* (pJFM41) | Wild-type *rpsL* insert | This study |
| pALO277:*rpsL*<sub>I82N</sub> (pJFM42) | Mutant *rpsL* insert | This study |
| pCas | Cas9, λ-Red and temperature-sensitive replicon | [31] |
| pTarget | sgRNA plasmid | [31] |
| pTarget:*rpsL* (pJFM40) | *rpsL*-specific sgRNA plasmid | This study |
| pSG3/4UGA | *lacZ* reporter containing a premature UGA codon | [26] |
| pSG12-6 | *lacZ* reporter containing a premature UAG codon | [26] |
| pOmpLac-M6 | Translational *ompA-lacZ* fusion with the M6 target sequence | [10] |
| pMicA-M6 | MicA-expression plasmid with the complementary M6 sequence | [10] |
| pControl | Promoterless control plasmid | [10] |
| pTP11 | Vector control for GcvB reporter assays | [12] |
| pPL-*gcvB* | GcvB-expression plasmid | [12] |
| pXG-10sf-*asd* | sfGFP translational fusion | [12] |
| pXG-10sf-*kgtP* | sfGFP translational fusion | [12] |
| pXG-10sf-*asc* | sfGFP translational fusion | [12] |

**Supplementary Table S5.**
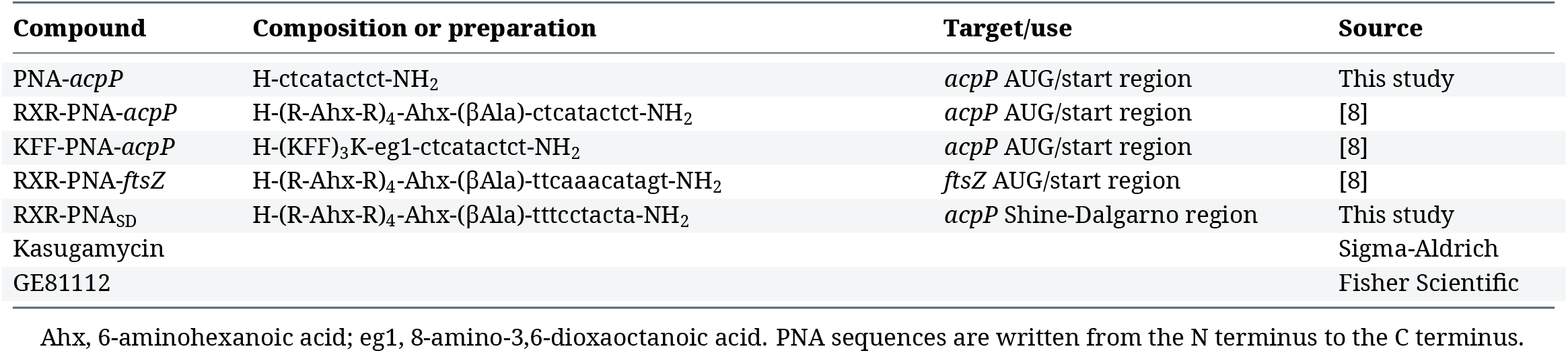
PNA and antimicrobial compounds.

**Supplementary Table S6.**
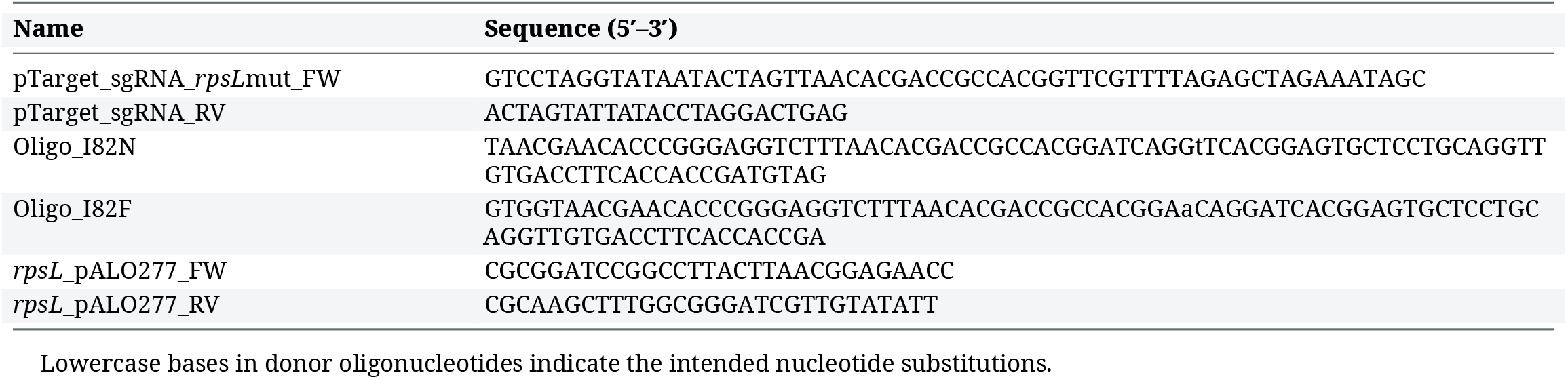
Oligonucleotides and genome-editing reagents.

